# How to choose sets of ancestry informative markers: A supervised feature selection approach

**DOI:** 10.1101/759464

**Authors:** Peter Pfaffelhuber, Franziska Grundner-Culemann, Veronika Lipphardt, Franz Baumdicker

**Author notes:** **Highlights:** - We provide AIMsetfinder, a tool to systematically select ancestry informative markers (AIMs). - Simulations of human population structure can be used to assess the performance of AIM selection procedures. - 17 SNPs identified by AIMsetfinder suffice to classify all african, european, east asian, and south asian individuals in the 1000 Genomes project correctly.

## Abstract

Inference of the Biogeographical Ancestry (BGA) of a person or trace relies on three ingredients: (1) A reference database of DNA samples including BGA information; (2) a statistical clustering method; (3) a set of loci which segregate dependent on geographical location, i.e. a set of so-called Ancestry Informative Markers (AIMs). We used the theory of feature selection from statistical learning in order to obtain AIM-sets for BGA inference. Using simulations, we show that this learning procedure works in various cases, and outperforms ad hoc methods, based on statistics like *F*_*ST*_ or informativeness for the choice of AIMs. Applying our method to data from the 1000 genomes project (excluding Admixed Americans) we identified an AIMset of 17 SNPs, which partly overlaps with existing ones. For continental BGA, the AIMset outperforms existing AIMsets on the 1000 genomes dataset, and gives a vanishing misclassification error.

## 1 Introduction

The biogeographical ancestry of a person (or trace) was coined in work leading in 2004 to the patent application US2004/0229231A1 by Toni Frudakis and Mark Shriver [17]. BGA has been defined as the *heritable component of “race” or heritage, which is relevant on any scale of resolution* [16]. Today, several application fields have formed, which range from population history (e.g. [12]), biomedicine (e.g. [19]) to personal genomics (e.g. [4]) and forensic genetics (e.g. [36]). In the present work, we are interested in the continental scale of BGA, and we step back from the above definition and either use the sampling location (in simulations) or the population origin, as given in a reference database such as the 1000 genomes dataset [1] as an equivalent of BGA.

In the forensic context, the available DNA trace is often only available in minimal amounts and might be contaminated with non-human, for instance microbial DNA. Thus inference of BGA is usually based on a set of only a few genetic markers rather than the whole genome. Therefore an important task is to find *Population-Specific Alleles* [48] or *Ancestry Informative Markers* (AIMs). Once such a set of markers (also called AIMsets below) is available, a classifier is used to group samples into BGA-classes. Here, we follow the frequently used approach to use a naïve Bayesian classifier, which has its roots in the software STRUCTURE [40], and has gained further interest through the software SNIPPER [38]; see also http://mathgene.usc.es/snipper/. It has recently been shown that for non-admixed samples, the naïve Bayesian approach outperforms logistic regression classifiers and genetic distance algorithms [8, 9].

AIMsets are often designed to distinguish specific populations. For example, the SNPforID 34-plex was introduced as a panel of 34 unlinked AIMs designed to distinguish between African (AFR), European (EUR), and East Asian (EAS) ancestry [38]. (It was later revised in order to include a SNP specific for East Asia; [15].) Furthermore, the Pacifiplex panel of 29 SNPs [46] was introduced to complement the SNPforID 34-plex in order to additionally distinguish Oceanic samples. A panel introduced in [27] consists of 55 AIMs and is designed to draw world-wide distinctions. To help locate AIMs, either putatively neutral mutations are used, or mutations subject to strong regional positive selection in the recent past, which create local adaptations [2]. Today, several AIMsets have been considered for different distinctions; see e.g. [39] for a review.

For BGA inference, finding good AIMsets is an important task. In Table 1, we collected example criteria how AIMs have been selected in practice. Most of these approaches must be considered empirical in the sense that the AIMs were not chosen in order to minimize the number of errors in a given task to infer BGA, but based on summary statistics such as *F*_*ST*_, the allele frequency differential, *δ*, or the informativeness, usually abbreviated *I*_*n*_ and introduced by [42] (see also in *Materials and Methods*). As explained in [36], roughly *F*_*ST*_ is proportional to 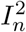, and similarly to the population frequency differential, such that ordering SNPs according to any of the above measures results in similar AIMsets.

**Table 1.** Example for empirical criteria of finding AIMsets.

|  |  |
| --- | --- |
| Sanchez et al, 2006 [44] | (i) not too low heterozygosity (i.e. at least 30%, which equals 0.28 for the minor allele frequency) in at least one population; and not too low heterozygosity (i.e. at least 20%, which equals 0.17 minor allele frequency) in all three populations; (ii) a minimum distance (100 kb) to neighboring genes; (iii) no likely association with commonly used STR loci; (iv) a small amplicon (less than 120 bp) generated for optimal primer design, as well as flanking DNA sequence reliably reported and free from interfering polymorphisms. |
| Phillips et al, 2007 [38] | (i) SNPs private to one or two populations, but non-segregating in others; (ii) an allele frequency differential between populations of $> 0.6$ between populations (the allele frequency differential is the absolute value of the difference in allele frequencies in two populations); (iii) tri-allelic SNPs; (iv) fixed difference markers, which separate one population (group) against other populations (or population groups). |
| Kosoy et al, 2009 [29]<br>and Nassir et al, 2009 [33] | (i) SNPs that overlapped with the 300K genome-wide Illumina SNP array; (ii) large allele frequency differences ($> 45\%$) between European and Amerindian groups and small allele frequency differences ($< 5\%$) between two disparate Amerindian groups; (iii) SNPs with large frequency differences ($> 45\%$) between African and European populations groups; (iv) minimum distance of AIMs (i.e. at least 8 cM on the deCODE map). |
| Kidd et al, 2014 [27] | (i) using AIMs found previously, e.g. in the AIMsets by [29] and [34]; (ii) SNPs with large pairwise allele frequency differences for a total of 63 populations; (iii) SNPs which perform well for classification using STRUCTURE [40]. |
| Phillips et al., 2014 [37] | (i) ranking SNPs by allele frequency differential; here, holding one population fixed, pairwise population comparison differentials were summed for the remaining populations; (ii) balancing the degree of differentiation among all population groups. |
| Jia et al, [24] | (i) only SNPs in Hardy-Weinberg equilibrium (HWE) were selected; (ii) population-specific markers, comprising private SNPs, or SNPs with a differential at least 0.5; (iii) an $F_{st}$-value of at least 0.35 (for three ancestral groups); (iv) no linkage disequilibrium (LD) among the population groups. |

A few algorithmic approaches in order to find AIMsets have also been given. While [27] uses STRUCTURE to assess the quality of AIMs, the use of Principal Component Analysis (PCA) and a clustering algorithm [35], or of eigenfunctions and support vector machines [55] in order to find an AIMset, were described. Conceptual approaches to find AIMsets have been given in [41]. Two of them add SNPs to a given AIMset one at a time. (i) The *univariate accumulation method* uses the SNPs which have the highest prediction accuracy when used as single SNPs. As discussed in [41], this method does not account for shared information between SNPs. While following such a univariate method, Zhao et al. add a penalty term to account for such shared information [56]. (ii) In the *greedy accumulation method*, the SNP with the highest prediction accuracy is used, when added to the current AIMset. We will use this greedy approach, and note that [43] follows a similar route. These authors provide an AIMset with 100 SNPs for distinguishing 24 populations (all from the HGDP dataset), and end up with an error rate of 40%. While being conceptually similar to [43] and [41], there are some important differences to our approach as detailed below. Most importantly we use a Bayesian prior distribution for BGA (rather than using a Maximum-Likelihood approach), resulting in a closed form for the estimate of allele frequencies, and we use logarithmic loss rather than misclassification error for choosing AIMs, which naturally allows the AIM selection procedure to account for the confidence of it’s predictions.

In inference methods for BGA and elsewhere, it is often assumed that alleles in different geographical origins come in different frequencies. Population genetic theory, in particular the so-called coalescent process (e.g. [52]), is an important tool for understanding the occurrence of such frequency differences. In addition, since its introduction by Kingman and Hudson [28, 23], it is also a tool for simulating genetic data for neutral variants. Note that the coalescent is versatile, and recombination, gene conversion and population size changes were subsequently included. Today, coalescent simulations play some role in association mapping of complex traits (see e.g. [57]), and form the foundation of various methods in population history (see e.g. [11]). Here, we use the coalescent for generating data in order to give a proof of principle that our approach for finding AIMsets works in independent simulated data. For this, we used an implementation from [26], which makes genome-wide simulation feasible.

In the language of statistical learning, inference of the BGA of a genetic trace is a high-dimensional classification problem. Here, the number of dimensions is the number of SNPs in the dataset. Finding AIMs in this context is then termed *feature selection* [20]. As such, general learning methods can be applied to the inference of BGA. To date, variants of STRUCTURE [40] are frequently used for the classification procedure. While the first application of this software was the unsupervised problem of finding structure in large datasets, reference databases incorporating individuals with (putatively) known ancestry make inference of BGA a supervised learning procedure. Hence, the approach has been taken further and software such as SNIPPER [38] has been developed. In the following, we subsume these approaches using the term *naïve Bayes* [32].

## 2 Materials and Methods

Before we come to our method how to choose AIMs in Section 2.2, we have to introduce some notions. Note that our implementation (which relies on python and R) can be downloaded from https://github.com/fbaumdicker/AIMsetfinder, and is described in detail in the SI.

### 2.1 Prerequisites

We assume that we have 2*n* + 2*m* (haploid) samples taken from a population, which is structured into *K* groups (or subpopulations), where *n*_*k*_ samples are from group *k* for *k* = 1, …, *K*. Using the *n* (diploid) samples (training set), we train a naïve Bayesian classifier (see below for the details). Once we have set up our classification method, we use the test set of (diploid) size *m* in order to obtain the misclassification error (also called test error; see below). Each sample consists of alleles at *s* SNPs (or loci). In sample *i* of deme *k* at SNP *j* we observe state *x*_*ijk*_ ∈ Ξ = {0, 1, 2}, which gives the number of copies of the derived allele in this sample. We stress that our method is independent of the knowledge of ancestral and derived alleles, such that *x*_*ijk*_ can as well indicate the number of copies of the less frequent allele. Moreover, since our method assumes linkage equilibrium between SNPs, we don’t need to know phase, i.e. the exact allelic compositions of chromosomes. Using these assumptions, we write *x*_*ijk*_ = *w*_2*i*−1,*j,k*_ + *w*_2*i,j,k*_, *i* = 1, …, *n*_*k*_, where *w*_2*i*−1,*j,k*_, *w*_2*i,j,k*_ ∈ {0, 1} are the two copies of sample *i* in SNP *j* in deme *k*. The frequency of the derived allele (or the less frequent allele) in deme *k* in SNP *j* is denoted *p*_*kj*_. Throughout, we will assume that we do not have three or four-fold degenerate sites in the sample.

#### Informativeness

Wright’s fixation index *F*_*ST*_ was introduced in order to have a measure for the amount of structure within a population [22]. Therefore, one can use SNPs with high values of *F*_*ST*_ to select AIMs. A more advantageous approaches is taken in [42], where a measure of global informativeness of a marker with respect to ancestry is introduced. Assume *K* islands and a bi-allelic marker with frequencies *p*_1_, …, *p*_*K*_ and 1 − *p*_1_, …, 1 − *p*_*K*_, respectively. Informativeness *I*_*n*_ is related to Shannon entropy, and defined as follows: For 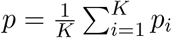,

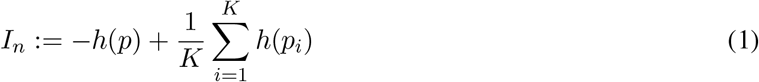

with *h*(*x*) ≔ *x* log(*x*) + (1 − *x*) log(1 − *x*) where we have the convention that *h*(0) = 0. We will use *I*_*n*_ both, for preselecting AIMs and to obtain a full set of AIMs; see below.

#### Naïve Bayesian classifier

For our Bayesian classifier, we implemented a simpler version of STRUCTURE [40]. Our aim is to classify diploid samples into *K* ≥ 2 groups using genetic data at *s* loci. Such data is described by (*yz*) ∈ Ξ^*s*^, by which we denote the number of copies of the derived (or less frequent) allele at all *s* loci. We use this notation since *y* and *z* can as well be seen as haploid genomes themselves, but only the number of copies is used in our algorithm. For a genome (*yz*) ∈ Ξ^*s*^, the classification method we use is based on probabilities *p*_*k*_(*y*) for a (haploid) sample *y* belonging to group *k*, which have to be obtained. Observing a genotype *yz* then gives the corresponding probabilities *p*_*k*_(*yz*) ≔ *p*_*k*_(*y*)*p*_*k*_(*z*)/(Σ_*ℓ*_*p*_*ℓ*_(*y*)*p*_*ℓ*_(*z*)) for the sample to belong to group *k*. (Here, we are actually assuming that all populations are in Hardy-Weinberg equilibrium.) Classification then is built upon the majority vote

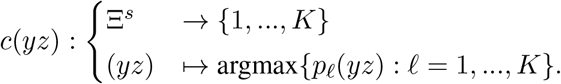

The values for *p*_*k*_(*y*) are obtained as follows: Let *w*_*ik*_ = (*w*_*i*1*k*_, …, *w*_*isk*_) be the SNPs of the *i*th (haploid) sample from the *k*th group in the training set, *i* = 1, …, 2*n*_*k*_, *k* = 1, …, *K*. For each SNP *j* in each of the *K* groups, the prior for the frequencies *q*_*jk*_ = (*q*_*jka*_)_*a*∈{0,1}_ are Dir((1)_*a*∈{0,1}_)-distributions, i.e. *q*_0_ = 1 − *q*_1_ is uniformly distributed on [0, 1]. Then – since the training data consists of a multinomial pick using the frequencies from the prior – standard theory [14] tells us that the posterior probability distribution of the allelic frequencies in group *k* are

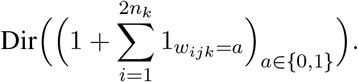

(Recall that if (*X*_*a*_)_*a*∈{0,1}_ ~ Dir((*x*_*a*_)_*a*∈{0,1}_) and *x* = *x*_*a*_, then 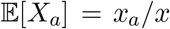 is the probability that a random pick with the random probabilities distributed according to the Dirichlet distributions is *a*, and 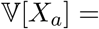 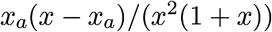, in particular the variance is smaller for higher *x*, i.e. larger sample sizes.) Using the naïve Bayes approach of assuming that all SNPs are independent, we get that

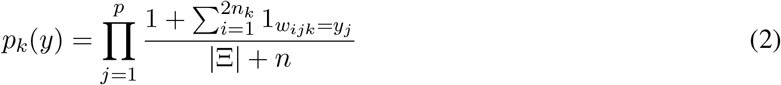

for the probability of picking a (haploid) sample *y* if we pick from group *k*. We note that SNIPPER slightly deviates from this formula, since (i) it uses an a priori probability for classification which is proportional to the population size in the reference dataset and (ii) it uses 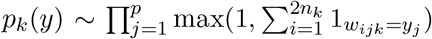 rather than(2). However, since 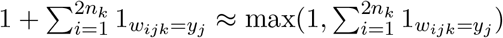 in large datasets, and (2) is derived directly using the Dirichlet prior, we stick to the approach described above. Note that adding 1 in the numerator of (2) is called Laplace smoothing in the naïve Bayes literature.

#### Misclassification error and logarithmic loss

If we have BGA information from all samples, the error we make in our classification scheme can be computed. We set *b*(*i*) = *k* if sample *i* was taken from population *k*. The misclassification (test) error is then, for the test data of size *m* with genotypes (*y*_*i*_*z*_*i*_)_*i*=1,…,*m*_,

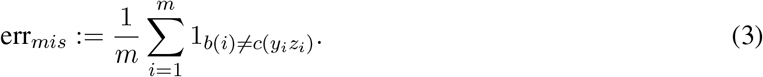

(But note that the misclassification error can as well be computed on the training set, which is then denoted the training error.) This error function does not take into account how confident the method is about its own belief, i.e. a small misclassification error can occur if *p*_*k*_(*y*_*i*_)*p*_*k*_(*z*_*i*_) is close to 1 for *b*(*i*) = *k*, or if *p*_*k*_(*y*_*i*_)*p*_*k*_(*z*_*i*_) is just marginally bigger than (*p*_*ℓ*_(*y*_*i*_)*p*_*ℓ*_(*z*_*i*_))_*ℓ*≠*k*_. This confidence is taken into account for a second error function we consider, the logarithmic loss (or logloss). It is defined as

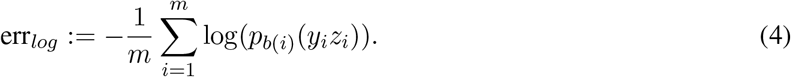

### 2.2 Feature selection: how to choose an AIMset

Most importantly, the Bayesian classifier as described above depends heavily on the selected *s* SNPs. We choose such an AIMset as follows, using only data on the training set: In order to reduce the number of possible AIMs, we compute the informativeness from (1) and exclude all SNPs with an informativeness below the 90% quantile (on the training set) of the resulting empirical distribution of informativenesses. We do this in order to improve the run-time of our algorithm without much performance deficit of the obtained AIMset. For the remaining SNPs, we start with an AIMset of size 0. For all SNPs, we determine the logarithmic loss (on the training set) when only this single SNP is used. The SNP with the smallest logloss is then put into the AIMset. Once we have an AIMset of size *s*, we try to add all remaining SNPs to this set and determine in each case the logloss (on the training set). Then, the (*s* + 1)st AIM is the SNP with the smallest logloss.

We found that minimizing the logloss (see (4)) for choosing AIMs works better than minimizing the misclassification error. The reason is that once the misclassification error is small, e.g. only two out of several thousand individuals are misclassified, the minimal misclassification error for the next AIM is not unique, and it is hard to come up with a rule which AIM should be chosen out of all minimizers. Since the logloss takes into account the exact frequencies at all AIMs, this minimum is nearly always unique, and no further rule is required.

Clearly, the step-wise approach we use here might not be optimal. The reason is that a global minimal logloss for *s* AIMs need not lie on a decreasing path of loglosses for 1, …, *s* AIMs. However, only this step-wise approach has a run-time which is linear in the size of the AIMset we are looking for. Given that real data frequently come with several million SNPs, this seems to be the only viable approach.

#### Cross-validation

Since the reported AIMs depend only on the genomic data on the training set, there is the chance that different AIMs appear on different training sets. In order to study this effect, we used five-fold cross-validation as follows: Splitting the data randomly in five parts of equal size, we take all five combinations of a training set comprising 80% of the individuals in the dataset and of a test set of the remaining 20% of the individuals. Then, finding AIMs step-wise as just described is carried out, leading to five different AIMsets for the five folds.

### 2.3 Simulated data

Population genetic data can efficiently be simulated by the coalescent [28, 23]. Today, a time-efficient, exact coalescent-simulator is available [26]. Such simulations result in data from (haploid) individuals, drawn uniformly from the continents (or islands or demes) of a structured population, which can then be combined to form diploids.

We run two sets of simulations. A detailed description is given in the SI. First, we use a symmetric migration model with five islands (mimicking five continents), i.e. migration rates between and sizes of islands are equal and constant over time. Here, migration rates (i.e. the expected number of haploids changing from one island to another island per generation) vary from 1 to 8. We simulated 20 pairs of chromosomes for 500 individuals on each island, where each recombining chromosome contains approximately 5 · 10^4^ SNPs. For obtaining an AIMset, we use two methods: (i) the 20 most informative SNPs (i.e. the SNPs with the highest informativeness, as given in (1)); (ii) the approach from Section 2.2.

Second, in order to be more realistic with respect to human history, we adopt the Out of Africa model as given in [18] for human genetic history. We stress that this model was recently implemented in https://github.com/popgensims/stdpopsim as a standard model for human population history. This model was fit using maximum likelihood on the allelic frequency spectrum of 5Mb of non-coding DNA (26,000 SNPs) from the Environmental Genome Project. Roughly, divergence between West African (as represented by 12 Yorubean from Ibadan, YRI) and Eurasian (represented by 22 individuals from a population from Utah, CEU) populations was inferred to have occurred 140 thousand years (kya) ago (the 95% confidence interval is 40 – 270 kya). The European and East Asian (represented by 12 Han Chinese, CHB) divergence time to was estimated to be 23 kya (95% confidence interval: 17 – 43 kya). In our simulations, we used the point estimators. Both Asian and European populations went through a bottleneck when they were founded. The most likely model also allows for recent migration between continents. Using this model, we simulate 20 pairs of chromosomes for 800 individuals in Africa, Asia and Europe. Again, the mutation rate is set such that we have about 5 · 10^4^ SNPs on each recombining chromosome. As in the symmetric case, we choose AIMsets either by taking the most informative SNPs, or by the approach as described in Section 2.2.

### 2.4 Human data

We downloaded 1000 Genomes data (phase 3) from ftp://ftp.1000genomes.ebi.ac.uk/vol1/ftp/release/20130502/, as well as information on the sampling locations from ftp://ftp.1000genomes.ebi.ac.uk/vol1/ftp/release/20130502/integrated_call_samples_v3.20130502.ALL.panel [1]. Note that this is data from 2504 individuals from five continental populations (called *superpops* in the dataset). These are 661 from Africa (AFR), 347 Admixed Americans (AMR), 504 East Asians (EAS), 503 Europeans (EUR) and 489 South-East Asians (SAS). The dataset comes with approximately 80 million SNPs. In the dataset, sampling locations are broken down to smaller populations, but we did not use this information and classified only according to the continental populations. Note that AMR individuals come from four different populations: Mexican Ancestry from Los Angeles (MXL); Puerto Ricans from Puerto Rico (PUR); Colombians from Medellin (CLM), Colombia; Peruvians from Lima, Peru (PUR). Also note that this dataset comes with no missing data.

#### Published AIMsets

For comparison of our method, we used published AIMsets from [33] (denoted the Seldin-AIMset), from [27] (denoted the Kidd-AIMset), and the EUROFORGENE global AIMset from [37] (denoted the EUROFORGENE-AIMset). We briefly introduce these AIMsets and their choice of markers:

- Seldin-AIMset: Validated on the HGDP dataset, this AIMset contains 93 AIMs and was designed to distinguish various populations from all continents (except Australia). Figure 2 in [33] indicates that 9 large population clusters were considered (two Africans, two Europeans, two South Asians, East Asia, Amerindian and Oceania). Wright’s *F*_*st*_ was used as a measure to differentiate between populations.
- Kidd-AIMset: Developed for forensic use, several approaches to select the 55 AIMs in this set were applied. Mainly, sites were included which have a large absolute frequency difference or large *F*_*st*_ values, where 63 populations with a total of 3071 individuals were tested (including the HGDP-data). The AIMset was tested against the Seldin AIMset and the AIMset from [34], both of which are based on HGDP-data.
- EUROFORGENE-AIMset: As stated in [37], this AIMset was chosen using an algorithm using just AIMs with a high sum of allele frequency differentials over population pairs, *δ*. This study selected AIMs also with respect published sources of AIMs. This resulted in an AIMset of 128 SNP, including 6 tri-allelic SNPs.

Note that the Seldin- and Kidd-AIMsets were combined to form the *Precision ID Ancestry panel*, which is also commercially available from ThermoFisher Scientific [51, 3].

## 3 Results

### 3.1 Simulated Data

#### Symmetric island model

When simulating data in a structured population with five islands, each pair sharing the same number of expected migrants per generation (the so-called migration rate), it is clear that the misclassification error (see (3)) increases with migration rate. Here, based on ~ 10^6^ SNPs we trained the classification method on a training set of 200 (diploid) individuals per island and then evaluated the test error on the remaining 50 individuals per island. When the classifier just uses the 20 most informative SNPs as AIMset, the set selected with the naïve Bayesian classifier that we implemented outperforms the AIMsets selected based on informativeness; see Figure 1.

**Figure 1.**
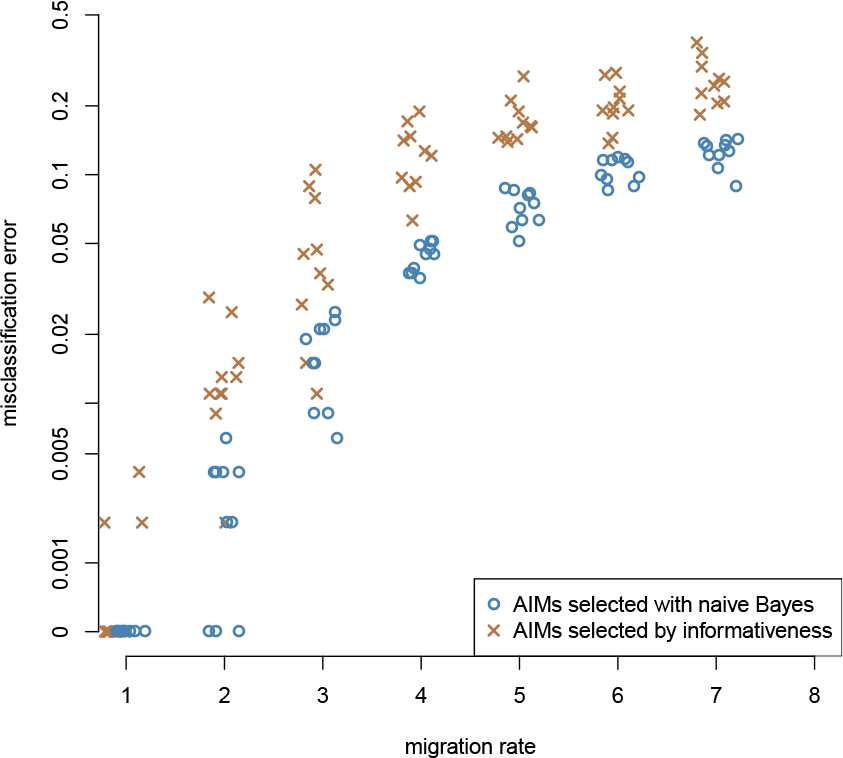
For varying migration rate *m* (which indicates the number of haploids migrating between any pair of islands in one generation), we obtain the misclassification error, when AIMs are selected using informativeness alone, or the naïve Bayes approach. In each run, we select 20 AIMs from a total of ~ 10^6^ SNPs.

**Figure 2.**
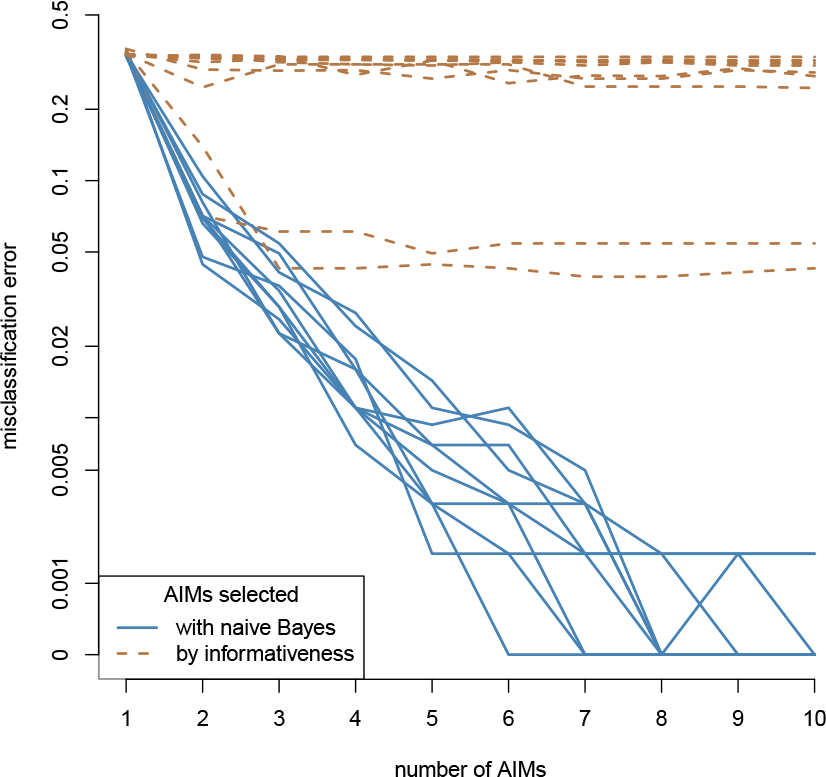
For a model for human evolution from [18], we determine the misclassification error depending on the number of AIMs used. Each line corresponds to one out of ten independent simulations.

#### Model for human evolution from [18]

The Out-of-Africa model from [18] aims at describing a realistic scenario for human evolution. As a proof of principle, we used a training set of 600 individuals in each continent (Europe, Africa, Asia), and a test set of 200 individuals in each continent. When using the most informative SNPs as AIMs, we see in Figure 2 that the misclassification error is never below 5%. The reason is the symmetry of the definition of informativeness, in conjunction with the asymmetric model for human evolution. However, when using our naïve Bayes approach, 8-10 AIMs suffice to get a vanishing misclassification error on the test set. Here, AIMs are ordered by the step-wise manner described in Section 2.2:

### 3.2 Human Data

#### Excluding admixed Americans

Using a specific (random) distinction in training and test set, we obtain an AIMset on the 1000 genomes data via the method described above. The goal of this set is a minimal error on the test set with a low number of AIMs, as described in *Materials and Methods*. At least when excluding Admixed Americans (AMR) from the dataset, we are able to achieve a vanishing misclassification (test) error (see (3)) with 21 AIMs; see Figure 3. See the Appendix for SNP identifiers. Interestingly, we did not find AIMs which can distinguish the continental origin when including AMRs; see Figure 4. (The SNPs found in this case are also in the Appendix.) We also note that the misclassification error when using the Kidd-AIMset, Seldin-AIMset or EUROFORGENE-AIMset is around 15%, since AMRs are not classified correctly. The reason might be that many AMRs have a mixed background and for this reason, we exclude admixed Americans in the sequel. Specifically, we follow [37], who also excluded the majority of AMRs for finding AIMs.

**Figure 3.**
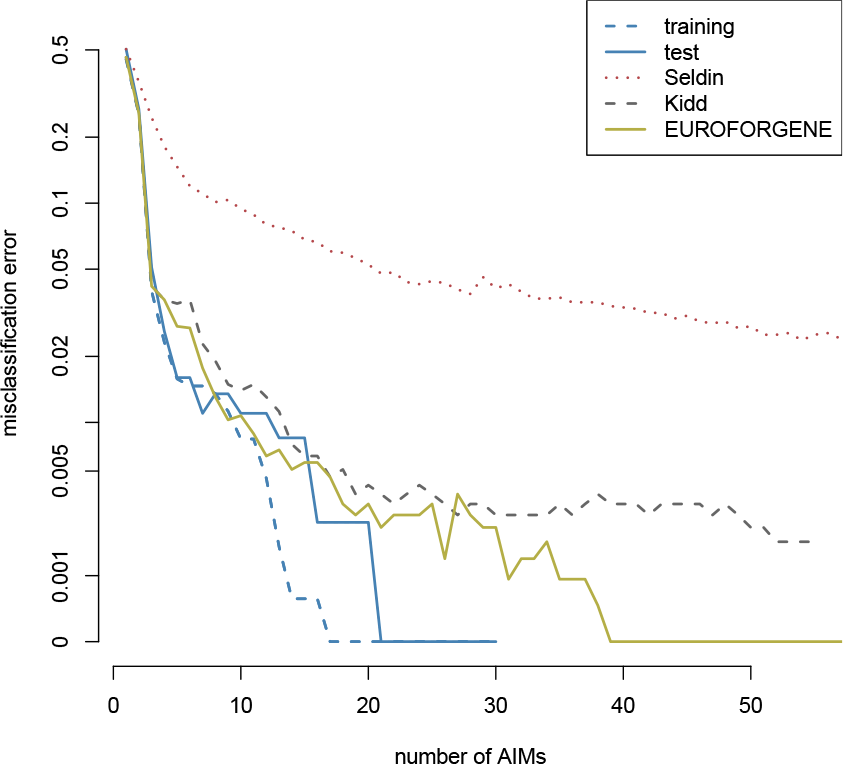
Comparison between misclassification errors on the 1000 genomes dataset (excluding AMRs). We only classified using continental origin of the sample.

**Figure 4.**
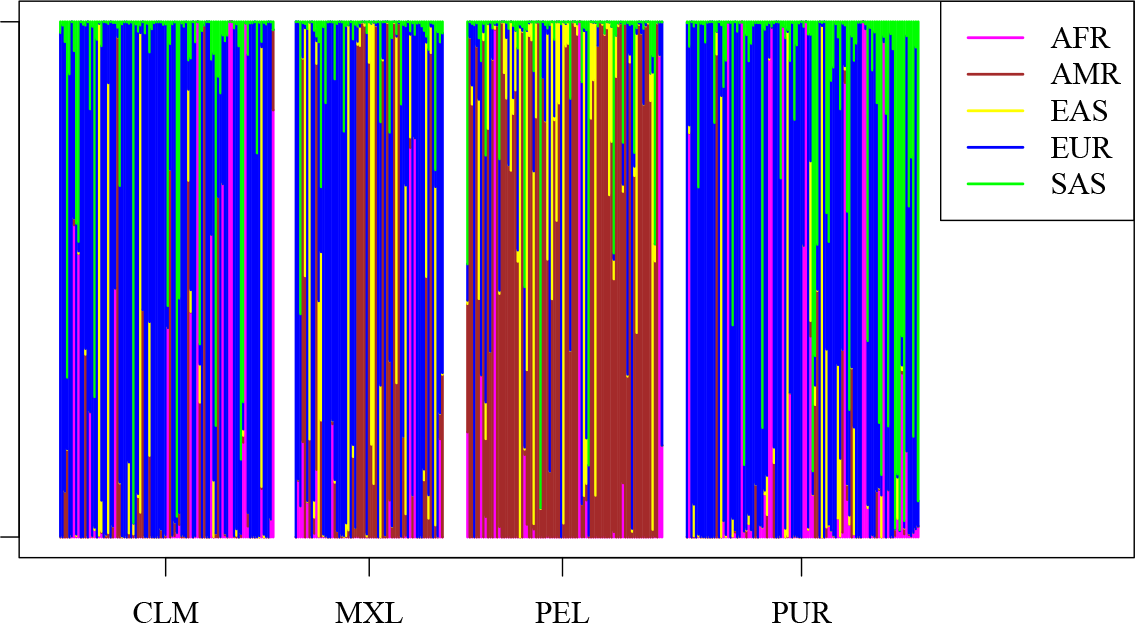
The classifier gives prediction probabilities from which continent a sample comes. When including AMRs in our algorithm, the AMR-samples are rarely classified into the class AMR. Rather, ancestry from other continents dominates the classification of AMRs.

#### Comparison to published AIMsets

After excluding AMRs, we compared the misclassification errors for continental origin (which drops to 0 at 21 AIMs) of our AIMset with published AIMsets by [33] (Seldin-AIMset), [27] (Kidd-AIMset), and [37] (EUROFORGENE-AIMset) on the 1000 genomes data. We have to mention that not all AIMs could be included for the EUROFORGENE AIMset, since (i) two AIMs are not included in the 1000 genomes dataset and (ii) not all are bi-allelic and our algorithm as it stands only works for bi-allelic SNPs. In total, we could use 121 (out of 128) AIMs in the EUROFORGENE AIMset; see Table 2 for details. In our analysis, the Seldi AIMset produces misclassification errors of approximately 3%, which is much higher than for the Kidd and EUROFORGENE AIMsets. The latter comes down to 0 at 40 AIMs; see Figure 3. Again, we ordered the AIMs appearing on the *x*-axis according to the minimal logloss (see (3)) an additional AIM produces in the naïve Bayes approach as described in *Materials and Methods*, i.e. by the same approach as we used for finding AIMs.

**Table 2.** ^0^ Informativeness was computed using the training set as in Figure 3. ^1^ SNP rs4471745 is not among the 10% most informative SNPs. ^2^ SNPs rs12402499 and rs17287498 (tri-allelic) are not present in the 1000 genomes dataset. ^3^ SNPs rs2069945, rs2184030, rs433342, rs4540055, rs5030240 are tri-allelic. ^4^ SNPs rs1509524, rs2139931, rs715605, rs9908046 are not among the 10% most informative SNPs.

|  | Kidd | Seldin | EUROFORGENE |
| --- | --- | --- | --- |
| total number of SNPs | 55 | 93 | 128 |
| in 1000 genomes dataset | 55 | 93 | 126<sup>2</sup> |
| ... and bi-allelic | 55 | 93 | 121<sup>3</sup> |
| ... and most informative<sup>0</sup> | 54<sup>1</sup> | 93 | 117<sup>4</sup> |

#### Cross-validation and remarks on some SNPs

Five-fold Cross-validation was used to see if the AIMset remains stable under changes in the training and test set. Here, for each of the five folds, we let our algorithm run until 10 AIMs were found. (The exact sequences of AIMs for the five folds are found in the Appendix.) Interestingly, among the first five AIMs in each fold, we only find eight different SNPs. Four of them also appear in both, the Kidd and EUROFORGENE AIMset:

- rs2814778: It is known that this SNP is part of the molecular basis of the Duffy blood group system [21] and relevant for malaria resistance [31]. It shows a nearly fixed difference between Europeans and Africans; see also Table 3.
- rs16891982: As part of the Irisplex system, which consists of six SNPs relevant for eye (or iris) color [54], this SNP is known to play a role in human pigmentation. It is also part of Hirisplex-S, a system consisting of 41 SNPs for simultaneous determining human eye, hair and skin color [53, 6].
- rs3827760: This SNP was recently found to be associated with eyebrow thickness [13] and facial morphology in Eurasians [30].
- rs1426654: Also part of the Hirisplex-S system [53, 6], this SNPs was shown to have a high correlation with skin and iris pigmentation in individuals of south Asian ancestry [49, 25].

**Table 3.** An AIMset with 17 SNPs which gives a perfect continental classification on the 1000 genomes dataset (if admixed Americans are excluded).

|  | rs2814778 | rs16891982 | rs3827760 | rs1872600 | rs1426654 | rs889603 | rs34454770 | rs5743618 | rs7541623 |
| --- | --- | --- | --- | --- | --- | --- | --- | --- | --- |
| AFR | 0.964 | 0.036 | 0.003 | 0.523 | 0.926 | 0.047 | 0.013 | 0.964 | 0.86 |
| EAS | 0 | 0.006 | 0.873 | 0.074 | 0.988 | 0.997 | 0.085 | 0.994 | 0.794 |
| EUR | 0.006 | 0.938 | 0.011 | 0.152 | 0.003 | 0.42 | 0.353 | 0.327 | 0.07 |
| SAS | 0 | 0.059 | 0.013 | 0.831 | 0.315 | 0.874 | 0.743 | 0.926 | 0.108 |
|  | rs114429736 | rs6053171 | rs10130486 | rs76057839 | rs2073917 | rs2847495 | rs1641127 | rs1355343 |  |
| AFR | 0.074 | 0.393 | 0.854 | 0.12 | 0.21 | 0.522 | 0.784 | 0.59 |  |
| EAS | 0.625 | 0.982 | 0.071 | 0.663 | 0.733 | 0.997 | 0.849 | 0.022 |  |
| EUR | 0.097 | 0.083 | 0.383 | 0.828 | 0.67 | 0.25 | 0.116 | 0.519 |  |
| SAS | 0.612 | 0.598 | 0.087 | 0.357 | 0.961 | 0.81 | 0.142 | 0.101 |  |

The other four (rs1872600, rs17034666, rs200750143, rs260687) have no associated publications in the dbSNP database [47].

#### Using the whole dataset for training (excluding AMRs)

Frequently, a difference between training and test error is attributed to overfitting the set of features (AIMset) to the training data. As can be seen from Figure 3, this is hardly the case here because the test error also drops to 0. Since the training and test error do not deviate too much, we used our approach to find an AIMset based on the whole dataset (but still excluding AMRs) for training. The resulting set is found in Table 3, where we also list allelic frequencies in the four classes (AFR=Africa, EAS=East Asia, EUR=Europe, SAS=South East Asia). Note that we again find the same overlap of SNPs with the Kidd and EUROFORGENE AIMset as above. We also note that the misclassification error when using only these four SNPs is 3.52% (when using the whole dataset but excluding admixed Americans as training and test-set). Notably, 14 of the selected SNPs fall within gene coding regions which highlights the efficiency of sites under selection for the inference of biogeographical ancestry. In addition, two of the 17 SNPs (rs2814778 and rs16891982) were found in a recently identified set of 27 SNPs that convey information on diverse health related Off-target phenotypes [5]. However, our method could readily be applied using a restricted subset of SNPs filtered by custom criteria.

#### Run-times

Cherry-picking AIMs without prior knowledge from a dataset as large as the 1000 genomes project with its approximately 80 million SNPs is a computationally demanding task. After filtering for only bi-allelic SNPs and highly informative SNPs (recall that we restrict ourselves to the top 10% for informativeness), approximately 8 million SNPs remain. In our step-wise manner for choosing the AIMset, we have to compute the logloss for all of these SNPs in order to decide which one to pick. When using a computer with 32 cores and not less than 64GB RAM, this task took (using multi-core support) about five hours to find one AIM, or about four days to find the AIMset from the last section. Note that the step-wise manner to choose AIMs results in run-times which are linear in the number of SNPs in the dataset.

## 4 Discussion

Using statistical learning, we obtained an automated method for producing an AIMset for biogeographical inference using genome-wide population data. As a core principle for finding the AIMset, we use an approach previously described by Rosenberg in [41], who calls the method *greedy* by the stepwise approach of adding SNPs to an AIMset. More specifically, we add SNPs one by one to the AIMset, which minimizes the logloss when classifying a training set using a naïve Bayesian classifier. In contrast to the misclassification error, the logloss accounts for the confidence in its classification being correct. In other words, choosing AIMs by minimizing the logloss leads to an AIMset which is more likely to classify traces with higher confidence. Once the AIMset is found, we again use the naïve Bayesian classifier for inference of BGA, as in [38].

A similar method (called AIM-SNPtag) for choosing an AIMset was introduced in [56], also based on the 1000 genomes dataset. First, combinations of classes/populations which will be most important to distinguish are manually determined. Second, for each of these combinations, SNPs (20.000) with a high *F*_*ST*_-value are pre-selected. Third, a feature selection algorithm (based on a naïve Bayesian classifier) is used in order to (i) minimize the misclassification error (for single SNPs) and (ii) minimize correlations between AIMs. However, since the error is only minimized on single SNPs in the last step, combinations of AIMs are not tested as in the stepwise approach we use here.

Although no manual step is involved in our approach, we are able to give an AIMset of 17 SNPs with a perfect classification of the 1000 genomes dataset into four continental groups (after excluding admixed Americans). It thereby outperforms other methods (which also exclude admixed Americans). However, we can still suggest some possible improvements of our method: (i) change the stepwise choice of the AIMs to an optimal choice of a combination of SNPs (called *exhaustive evaluation* in [41]), but note that the computational complexity is huge; (ii) allow for deviations from independence of the two copies for each locus (Hardy-Weinberg equilibrium) within populations, as is done in the extension of SNIPPER [45]; see [8, Online resource 1]; (iii) rather than treating all SNPs the same in (2), more important SNPs should get a larger weight in the actual classification procedure; (iv) rather than only using bi-allelic SNPs, also tri-allelic SNPs or microhaplotypes could be used [7].

For comparing the performance of our 17-SNPs-AIMset to published AIMsets (Kidd [27], Seldin [33], EUROFORGENE [37]), we excluded Admixed Americans from the 1000 genomes dataset since the naïve Bayesian classifier was not able to classify them by any AIMset. The Seldin AIMset (93 SNPs) was announced to be able to distinguish between the continental groups of Oceana, South Asia, East Asia, Sub-Saharan Africa, North and South America, and Europe, and was tested on HGDP, HapMap and other data. Our results show that this AIMset is outperformed by the smaller Kidd AIMset (55 SNPs) for inference of continental BGA (African, European, East Asian, South-East Asian) on the 1000 genomes dataset. The Kidd AIMset in turn was not intended to classify according to continents, but according to 63 populations. The EUROFORGENE AIMset was designed to draw distinctions between Africans, Europeans, East Asians, Native Americans and Oceanians, but not South Asians. Still, this AIMset leads to a vanishing misclassification error on the 1000 genomes dataset with the right subset of 40 AIMs. While our AIMset is able to perfectly classify the 1000 genomes dataset with as little as 17 SNPs, it was not trained for any other distinction.

In [27], it was conjectured that “the greatest improvement [in BGA inference] will come from using *better* SNPs. The problem is finding SNPs that provide a clearer differentiation of certain populations or groups of populations without detracting from differentiation among some other populations.” Here, using the 1000 genomes dataset, only some SNPs appear simultaneously in several AIMsets (rs2814778, rs16891982, rs3827760, rs1426654, which appear in the Kidd, EUROFORGENE and our AIMset). As our cross-validation results show, the remaining AIMs depend not only on the populations which shall be distinguished, but also on the individuals present in the training and test set. Therefore, we conclude that for a given classification task there exists a multitude of sets with comparable performance to the optimal AIMset. However, the performance of all AIMset and methods is worse when admixed Americans are included in training and test set.

Simulation techniques provide benefits in various respects. In population genetics, powerful software based on the coalescent [23] is today available which can simulate genome-wide data [26]. Once a population model is specified, it generates genealogical relationships at all loci under consideration. Furthermore, once a population model is specified, simulations come with the advantage of “perfect” sampling: BGA in simulations can only mean the simulated sampling location when running the software. In this case, “perfect” sampling means that the technique recruits simulated individuals only on the basis of their simulated belonging to one of the populations designated in the previously defined classification scheme and no ambiguous cases can occur.

We nevertheless believe that simulation techniques can furthermore be beneficial in forensic genetics, and provide a proof of principle in scenarios including: (i) comparison of misclassification errors for autosomal (recombining) and mitochondrial- or Y-chromosomal (non-recombining) data; (ii) determine the performance of classifiers if tested data comes from populations that are missing in the reference database; (iii) study the spatial resolution of classification of BGA with a given number of AIMs; (iv) evaluation of classifiers for admixed populations.

The disadvantage of simulated data is the impossibility to simulate a real-life scenario of human population history since all models require simplification. For example, migration rates dramatically increased in the past decades, but the model for human evolution used in our simulation study from [18] was fitted using only Yorubeans, Europeans from Utah and Han Chinese individuals. Furthermore, populations are not always and not everywhere so neatly delimitable from each other.

The method we propose to find AIMsets starts off with a reference dataset. An apparent complication in the 1000 genome dataset is the group of Admixed Americans, including individuals coming from Mexico, Puerto Rico, Colombia and Peru. “Admixed Americans” then seems to relate to the US census category “Hispanics”. As we have seen, these are hard to classify into a single continental group, but were nevertheless kept and labelled as a separate group when designing the 1000 genomes dataset. Still, excluding admixed individuals and populations from reference databases in both, training- and test-sets, is a common practise [37], [56] and makes our method work perfectly, but it may not be working well when applied in a social reality in which admixed individuals make up a considerable proportion of the population (e.g. in many urban areas worldwide, and in postcolonial societies).

Another complication of inference of BGA in practise lies in the genesis of reference datasets. Here, sampling decisions come with various constraints. Sampling for a reference database that aims at distinguishing populations needs to make assumptions that may not be congruent with social reality. Usually, reference databases depend on information about recent ancestry, provided by the test subject, and use it a proxy for BGA. In order to obtain reference samples from individuals that are most likely to have all of their ancestry in only one location or population, the information sought after is the origin of all four grandparents ([10, 12] and [50, p. 104]). This information, however, can be ambiguous, wrong, or based on different categories than the reference database’s classification provides. This points to the complexities that arise when genetic and social classification schemes do not map onto each other. Moreover, legal restrictions might protect some countries to be included in reference datasets, and socially loaded group-terms might bias the decision of individuals if they are asked to give their self-declared BGA.

A result of such sampling decisions becomes clear after a closer look at the 1000 genomes dataset. As an example, Europe is represented by five populations (Utah Residents with European Ancestry, Toscani in Italia, Finns, British, and the Iberian Population in Spain). According to the Informed Consent sheet of the 1000 genomes project [10], “the samples should be from people whose grandparents mostly came from [geographic location or ethnic group]” (p. 2). This means that e.g. “British” does not refer to British citizenship or self-declared Britishness, but to individuals who have British ancestors, which is a subset of “British citizens”). Although our AIMset of 17 SNPs gives a perfect classification on this dataset, it is hard to predict the performance of the same AIMset on data sampled under these conditions from other parts of Europe, or for samples without any constraint on ancestry.

To sum up, BGA categories might frequently introduce slippages between social and genetic population labels, which makes the ultimate goal of BGA inference even more challenging: to provide a useful tool in forensic casework. After all, the task for forensic geneticists and investigators is to translate a population label from a genetic reference database into a social category helpful enough for focussing investigations.

## Acknowledgments

We thank Fabian Freund for useful hints on some statistical methods, as well as Jenny Reardon, Bernhard Haubold and Fabian Staubach for fruitful discussions. Thorsten Schmidt kindly provided important computational power for this project.

## Appendix

## A Human data

## Excluding admixed Americans

For the specific (random) distinction in training and test set, the AIMset of 30 SNPs is as follows (ordered in a step-wise manner for minimizing the logloss as described in the main text:
rs2814778, rs16891982, rs3827760, rs1872600, rs1426654, rs3768641, rs5743618, rs112790372, rs889603, rs6053171, rs7541623, rs1906496, rs2000639, rs1550778, rs260687, rs4587880, rs2654201, rs6060652, rs191090, rs7686018, rs12916300, rs1811510, rs12536329, rs1630897, rs10130486, rs1015594, rs2196051, rs58827274, rs200450082, rs9762896.

## Including admixed Americans

When we include admixed Americans (AMR in the dataset), the misclassification error becomes significant; see Figure 4. The resulting AIMs are here rs2814778, rs16891982, rs3827760, rs4900908, rs1133238, rs7152177, rs5750993, rs12091805, rs184261752, rs72818358, rs509360, rs11772526, rs2357234, rs12052770, rs10792010, rs145327787, rs62419475, rs117787287, rs8079722, rs228092, rs9788726, rs12916300, rs184261752, rs2789823, rs12221394, rs2597305, rs2357234, rs509360, rs9320800, rs9550774.

## Five fold cross-validation

When training the classifier on different training sets, different AIMsets result. We studied this phenomenon using five fold cross-validation. Here are the AIMsets (of size 10) we found in the five folds (ordered by the step-wise manner as described in *Materials and Methods*:

- Fold 1: rs2814778, rs16891982, rs17034666, rs200750143, rs1426654, rs889603, rs34454770, rs2654201, rs6053171, rs114429736.
- Fold 2: rs2814778, rs16891982, rs3827760, rs260687, rs200750143, rs3768641, rs6053171, rs2402522, rs76057839, rs114429736.
- Fold 3: rs2814778, rs16891982, rs3827760, rs200750143, rs1426654, rs13011477, rs12929243, rs998401, rs1058119, rs2654201.
- Fold 4: rs2814778, rs16891982, rs3827760, rs1872600, rs1426654, rs889603, rs2443857, rs6053171, rs112790372, rs371618232.
- Fold 5: rs2814778, rs16891982, rs3827760, rs1872600, rs1426654, rs889603, rs34454770, rs509360, rs10494991, rs1357368.

## References

[1] 1000 Genomes Project Consortium, Adam Auton, Lisa D. Brooks, Richard M. Durbin, Erik P. Garrison, Hyun Min Kang, Jan O. Korbel, Jonathan L. Marchini, Shane McCarthy, Gil A. McVean, and Gonçalo R. Abecasis. A global reference for human genetic variation. Nature, 526(7571):68–74, 2015.

[2] Joshua M. Akey, Michael A. Eberle, Mark J. Rieder, Christopher S. Carlson, Mark D. Shriver, Deborah A. Nickerson, and Leonid Kruglyak. Population history and natural selection shape patterns of genetic variation in 132 genes. PLoS Biology, 2(10):e286, 2004.

[3] Muna Al-Asfi, Dennis McNevin, Bhavik Mehta, Daniel Power, Michelle E. Gahan, and Runa Daniel. Assessment of the precision id ancestry panel. International Journal of Legal Medicine, 132(6):1581–1594, 2018.

[4] Misha Angrist. Personal genomics: Where are we now? Applied & Translational Genomics, 8:1–3, 2016.

[5] Cedric Bradbury, Anna Köttgen, and Fabian Staubach. Off-target phenotypes in forensic DNA phenotyping and biogeographic ancestry inference: A resource Forensic Science International: Genetics, 38:93–104, 2019.

[6] Lakshmi Chaitanya, Krystal Breslin, Sofia Zuñiga, Laura Wirken, Ewelina Pospiech, Magdalena Kukla-Bartoszek, Titia Sijen, Peter de Knijff, Fan Liu, Wojciech Branicki, Manfred Kayser, and Susan Walsh. The HIrisPlex-S system for eye, hair and skin colour prediction from DNA: Introduction and forensic developmental validation. Forensic Science International. Genetics, 35:123–135, 2018.

[7] E. Cheung, C. Phillipps, M. Eduardoff, M. Victoria Lareu, and D. McNevin. Performance of ancestry-informative SNP and microhaplotype markers. Forensic Science International. Genetics, (in press):1–10, 2019.

[8] E. Y. Y. Cheung, M. E. Gahan, and D. McNevin. Prediction of biogeographical ancestry from genotype: a comparison of classifiers. International Journal of Legal Medicine, 131:901–912, 2017.

[9] Elaine Y. Y. Cheung, Michelle Elizabeth Gahan, and Dennis McNevin. Prediction of biogeographical ancestry in admixed individuals. Forensic Science International. Genetics, 36:104–111, 2018.

[10] 1000 Genomes Project Consortium. 1000 genomes project: Developing a research resource for studies of human genetic variation. consent to participate. https://www.internationalgenome.org/sites/1000genomes.org/files/docs/Informed%20Consent%20Form%20Template.pdf, download 15.8.2019.

[11] A. J. Drummond, A. Rambaut, B. Shapiro, and O. G. Pybus. Bayesian coalescent inference of past population dynamics from molecular sequences. Molecular biology and evolution, 22(5):1185–1192, 2005.

[12] Eran Elhaik, Tatiana Tatarinova, Dmitri Chebotarev, Ignazio S. Piras, Carla Maria Calò, Antonella De Montis, Manuela Atzori, Monica Marini, Sergio Tofanelli, Paolo Francalacci, Luca Pagani, Chris Tyler-Smith, Yali Xue, Francesco Cucca, Theodore G. Schurr, Jill B. Gaieski, Carlalynne Melendez, Miguel G. Vilar, Amanda C. Owings, Rocío Gómez, Luca Pagani, Chris Tyler-Smith, Yali Xue, Francesco Cucca, Theodore G. Schurr, Jill B. Gaieski, Carlalynne Melendez, Miguel G. Vilar, Amanda C. Owings, Rocío Gómez, Ricardo Fujita, Fabrício R. Santos, David Comas, Oleg Balanovsky, Elena Balanovska, Pierre Zalloua, Himla Soodyall, Ramasamy Pitchappan, Arunkumar Ganeshprasad, Michael Hammer, Lisa Matisoo-Smith, R. Spencer Wells, and Genographic Consortium. Geographic population structure analysis of worldwide human populations infers their biogeographical origins. Nature Communications, 5:3513, 2014.

[13] Chihiro Endo, Todd A. Johnson, Ryoko Morino, Kazuyuki Nakazono, Shigeo Kamitsuji, Masanori Akita, Maiko Kawajiri, Tatsuya Yamasaki, Azusa Kami, Yuria Hoshi, Asami Tada, Kenichi Ishikawa, Maaya Hine, Miki Kobayashi, Nami Kurume, Yuichiro Tsunemi, Naoyuki Kamatani, and Makoto Kawashima. Genome-wide association study in japanese females identifies fifteen novel skin-related trait associations. Scientific Reports, 8(1):8974, 2018.

[14] D. Fink. A compendium of conjugate priors in progress report: Extension and enhancement of methods for setting data quality objectives. Tech. Rep., Montana State University, 1995.

[15] M. Fondevila, C. Phillips, C. Santos, A. Freire Aradas, P. M. Vallone, J. M. Butler, M. V. Lareu, and A. Carracedo. Revision of the SNPforID 34-plex forensic ancestry test: Assay enhancements, standard reference sample genotypes and extended population studies. Forensic Science International. Genetics, 7(1):63–74, 2013.

[16] T. N. Frudakis and M. D. Shriver. Compositions and methods for inferring ancestry, 2004. https://patentimages.storage.googleapis.com/dd/3c/d7/75365f60149c53/US20040229231A1.pdf, US Patent 0229231 A1.

[17] Lisa Gannett. Biogeographical ancestry and race. Studies in History and Philosophy of Biological and Biomedical Sciences, 47 Part A:173–184, 2014.

[18] Ryan N. Gutenkunst, Ryan D. Hernandez, Scott H. Williamson, and Carlos D. Bustamante. Inferring the joint demographic history of multiple populations from multidimensional SNP frequency data. PLoS Genetics, 5(10):e1000695, 2009.

[19] Indrani Halder, Kevin E. Kip, Suresh R. Mulukutla, Aryan N. Aiyer, Oscar C. Marroquin, Gordon S. Huggins, and Steven E. Reis. Biogeographic ancestry, self-identified race, and admixture-phenotype associations in the Heart SCORE Study. American Journal of Epidemiology, 176(2):146–155, 2012.

[20] T. Hastie, R. Tibshirani, and J. Friedman. The Elements of Statistical Learning. Springer, 2nd edition, 2008.

[21] Gabriela Höher, Marilu Fiegenbaum, and Silvana Almeida. Molecular basis of the Duffy blood group system. Blood Transfusion = Trasfusione del Sangue, 16(1):93–100, 2018.

[22] R. R. Hudson, M. Slatkin, and W. P. Maddison. Estimation of levels of gene flow from DNA sequence data. Genetics, 132(2):583–589, 1992.

[23] R.R. Hudson. Properties of a neutral allele model with intragenic recombination. Theo. Pop. Biol., 23:183–201, 1983.

[24] Jing Jia, Yi-Liang Wei, Cui-Jiao Qin, Lan Hu, Li-Hua Wan, and Cai-Xia Li. Developing a novel panel of genome-wide ancestry informative markers for bio-geographical ancestry estimates. Forensic Science International. Genetics, 8(1):187–194, 2014.

[25] M. Jonnalagadda, M. Faizan, S. Ozarkar, R. Ashma, S. Kulkarni, H. Norton, and E. Parra. A Genome-Wide Association Study of Skin and Iris Pigmentation among Individuals of South Asian Ancestry. Genome Biology and Evolution, 11:1066–1076, 2019.

[26] Jerome Kelleher, Alison M. Etheridge, and Gilean McVean. Efficient coalescent simulation and genealogical analysis for large sample sizes. PLoS Computational Biology, 12(5):e1004842, 2016.

[27] Kenneth K. Kidd, William C. Speed, Andrew J. Pakstis, Manohar R. Furtado, Rixun Fang, Abeer Madbouly, Martin Maiers, Mridu Middha, Françoise R. Friedlaender, and Judith R. Kidd. Progress toward an efficient panel of SNPs for ancestry inference. Forensic Science International. Genetics, 10:23–32, 2014.

[28] J.F.C. Kingman. The coalescent. Stochastic Process. Appl., 13(3):235–248, 1982.

[29] Roman Kosoy, Rami Nassir, Chao Tian, Phoebe A. White, Lesley M. Butler, Gabriel Silva, Rick Kittles, Marta E. Alarcon-Riquelme, Peter K. Gregersen, John W. Belmont, Francisco M. De La Vega, and Michael F. Seldin. Ancestry informative marker sets for determining continental origin and admixture proportions in common populations in america. Human Mutation, 30(1):69–78, 2009.

[30] Y. Li, W. Zhao, D. Li, X. Tao, Z. Xiong, J. Liu, W. Zhang, A. Ji, K. Tang, F. Liu, and C. Li. EDAR, LYPLAL1, PRDM16, PAX3, DKK1, TNFSF12, CACNA2D3, and SUPT3H gene variants influence facial morphology in a Eurasian population. Human Genetics, 138:681–689, 2019.

[31] Kimberly F. McManus, Angela M. Taravella, Brenna M. Henn, Carlos D. Bustamante, Martin Sikora, and Omar E. Cornejo. Population genetic analysis of the DARC locus (Duffy) reveals adaptation from standing variation associated with malaria resistance in humans. PLoS Genetics, 13(3):e1006560, 2017.

[32] K. Murphy. Naive Bayes classifiers. Technical Report, 2006. https://datajobs.com/data-science-repo/Naive-Bayes-[Kevin-Murphy].pdf.

[33] Rami Nassir, Roman Kosoy, Chao Tian, Phoebe A. White, Lesley M. Butler, Gabriel Silva, Rick Kittles, Marta E. Alarcon-Riquelme, Peter K. Gregersen, John W. Belmont, Francisco M. De La Vega, and Michael F. Seldin. An ancestry informative marker set for determining continental origin: validation and extension using human genome diversity panels. BMC Genetics, 10:39, 2009.

[34] Caroline M. Nievergelt, Adam X. Maihofer, Tatyana Shekhtman, Ondrej Libiger, Xudong Wang, Kenneth K. Kidd, and Judith R. Kidd. Inference of human continental origin and admixture proportions using a highly discriminative ancestry informative 41-SNP panel. Investigative Genetics, 4(1):13, 2013.

[35] Peristera Paschou, Elad Ziv, Esteban G. Burchard, Shweta Choudhry, William Rodriguez-Cintron, Michael W. Mahoney, and Petros Drineas. Pca-correlated snps for structure identification in worldwide human populations. PLoS Genetics, 3(9):1672–1686, 2007.

[36] C. Phillips. Forensic genetic analysis of bio-geographical ancestry. Forensic Science International. Genetics, 18:49–65, 2015.

[37] C. Phillips, W. Parson, B. Lundsberg, C. Santos, A. Freire-Aradas, M. Torres, M. Eduardoff, C. Børsting, P. Johansen, M. Fondevila, N. Morling, P. Schneider, EUROFORGEN-NoE Consortium, A. Carracedo, and M. V. Lareu. Building a forensic ancestry panel from the ground up: The EUROFORGEN Global AIM-SNP set. Forensic Science International. Genetics, 11:13–25, 2014.

[38] C. Phillips, A. Salas, J. J. Sánchez, M. Fondevila, A. Gómez-Tato, J. Álvarez Dios, M. Calaza, M. Casares de Cal, D. Ballard, M. V. Lareu, A.. Carracedo, and The SNPforID Consortium. Inferring ancestral origin using a single multiplex assay of ancestry-informative marker SNPs. Forensic Science International. Genetics, 1:273–280, 2007.

[39] Chris Phillips, Carla Santos, Manuel Fondevila, Ángel Carracedo, and Maria Victoria Lareu. Inference of Ancestry in Forensic Analysis I: Autosomal Ancestry-Informative Marker Sets. In Forensic DNA Typing Protocols, volume 1420 of Methods in Molecular Biology, pages 233–253. Springer, New York, 2016.

[40] J. Pritchard, M. Stephens, and P. Donnelly. Inference of population structure using multilocus genotype data. Genetics, 155:945–954, 2000.

[41] Noah A. Rosenberg. Algorithms for selecting informative marker panels for population assignment. Journal of Computational Biology: A Journal of Computational Molecular Cell Biology, 12(9):1183–1201, 2005.

[42] Noah A. Rosenberg, Lei M. Li, Ryk Ward, and Jonathan K. Pritchard. Informativeness of genetic markers for inference of ancestry. American Journal of Human Genetics, 73(6):1402–1422, 2003.

[43] Joshua N. Sampson, Kenneth K. Kidd, Judith R. Kidd, and Hongyu Zhao. Selecting SNPs to identify ancestry. Annals of Human Genetics, 75(4):539–553, 2011.

[44] Juan J. Sanchez, Chris Phillips, Claus Børsting, Kinga Balogh, Magdalena Bogus, Manuel Fondev-ila, Cheryl D. Harrison, Esther Musgrave-Brown, Antonio Salas, Denise Syndercombe-Court, Peter M. Schneider, Angel Carracedo, and Niels Morling. A multiplex assay with 52 single nucleotide polymorphisms for human identification. Electrophoresis, 27(9):1713–1724, 2006.

[45] C. Santos, C. Phillips, A. Gomez-Tato, J. Alvarez-Dios, A. Carracedo, and M. V. Lareu. Inference of ancestry in forensic analysis II: Analysis of genetic data. Methods Mol. Biol., 1420:255–285, 2016.

[46] Carla Santos, Christopher Phillips, Manuel Fondevila, Runa Daniel, Roland A. H. van Oorschot, Esteban G. Burchard, Moses S. Schanfield, Luis Souto, Jolame Uacyisrael, Marc Via, Ángel Carracedo, and Maria V. Lareu. Pacifiplex: an ancestry-informative SNP panel centred on Australia and the Pacific region. Forensic Science International. Genetics, 20:71–80, 2016.

[47] S. T. Sherry, M. H. Ward, M. Kholodov, J. Baker, L. Phan, E. M. Smigielski, and K. Sirotkin. dbSNP: the NCBI database of genetic variation. Nucleic Acids Research, 29(1):308–311, 2001.

[48] M. D. Shriver, M. W. Smith, L. Jin, A. Marcini, J. M. Akey, R. Deka, and R. E. Ferrell. Ethnic-affiliation estimation by use of population-specific DNA markers. American Journal of Human Genetics, 60(4):957–964, 1997.

[49] R. Stokowskia, P.V. Krishna Pant, T. Dadd, A. Fereday, D. Hinds, C. Jarman, W. Filsell, R. Ginger, M. Green, F. J. van der Ouderaa, and D. R. Cox. A Genomewide Association Study of Skin Pigmentation in a South Asian Population. American Journal of Human Genetics, 81:1119–1132, 2007.

[50] M. Stoneking. An introduction to molecular anthropology. Wiley, New York, 2017.

[51] ThermoFisher. Precision ID Ancestry Panel, 2016. https://www.thermofisher.com/content/dam/LifeTech/Document, download 8.8.2019.

[52] J. Wakeley. Coalescent Theory: An Introduction. Roberts & Company, 2008.

[53] Susan Walsh, Lakshmi Chaitanya, Krystal Breslin, Charanya Muralidharan, Agnieszka Bronikowska, Ewelina Pospiech, Julia Koller, Leda Kovatsi, Andreas Wollstein, Wojciech Branicki, Fan Liu, and Manfred Kayser. Global skin colour prediction from DNA. Human Genetics, 136(7):847–863, 2017.

[54] Susan Walsh, Fan Liu, Kaye N. Ballantyne, Mannis van Oven, Oscar Lao, and Manfred Kayser. Irisplex: a sensitive DNA tool for accurate prediction of blue and brown eye colour in the absence of ancestry information. Forensic Science International. Genetics, 5(3):170–180, 2011.

[55] Jun Zhang. Ancestral informative marker selection and population structure visualization using sparse Laplacian eigenfunctions. PloS one, 5(11):e13734, 2010.

[56] Shilei Zhao, Cheng-Min Shi, Liang Ma, Qi Liu, Yongming Liu, Fuquan Wu, Lianjiang Chi, and Hua Chen. AIM-SNPtag: A computationally efficient approach for developing ancestry-informative SNP panels. Forensic Science International. Genetics, 38:245–253, 2019.

[57] Sebastian Zöllner and Jonathan K. Pritchard. Coalescent-based association mapping and fine mapping of complex trait loci. Genetics, 169(2):1071–1092, 2005.

